# Retigabine Suppresses Loss of Force in a Mouse Model of Hypokalemic Periodic Paralysis

**DOI:** 10.1101/2022.05.20.492877

**Authors:** Marbella Quiñonez, Stephen C. Cannon

**Author notes:** Correspondence: Dr. Stephen Cannon, Department of Physiology, David Geffen School of Medicine at UCLA, 650 Charles E Young Dr Box 951751, Los Angeles, CA 90095-1751, USA.

## Abstract

**Objective:** The goal of this experimental study was to test the hypothesis that the potassium channel opener retigabine can prevent the episodic loss of force in hypokalemic periodic paralysis (HypoPP).

**Methods:** A knock-in mutant mouse model of HypoPP (*Scn4a* p.R669H) was used to determine whether pretreatment with retigabine suppressed the loss of force, or post-treatment hastened recovery of force for a low-K^+^ challenge in an ex vivo contraction assay.

**Results:** Retigabine completely prevents the loss of force induced by a 2 mM K^+^ challenge (protection) in our mouse model of HypoPP, with a 50% inhibitory concentration (IC_50_) of 0.8 µM. In comparison, the effective concentration for the K_ATP_ channel opener pinacidil was ten-fold higher. Application of retigabine also reversed the loss of force (rescue) for HypoPP muscle maintained in 2 mM K^+^.

**Interpretation:** Retigabine, a selective agonist of the K_V_7 family of potassium channels, is effective for the prevention of low-K^+^ induced attacks of weakness and to enhance recovery from an on-going loss of force in a mouse model of HypoPP. Substantial protection from the loss of force occurred in the low micromolar range, well within the therapeutic window for retigabine.

## INTRODUCTION

Hypokalemic periodic paralysis (HypoPP) presents with recurrent episodes weakness, often in association with low serum potassium ([K^+^] < 3.5 mEq/l)^1, 2^. Attacks of weakness are variable, both amongst affected members in a family or for a single individual over time, with regard to affected muscle groups, severity, frequency and duration. Trigger factors that increase the risk of experiencing an attack of weakness are a prominent feature in all forms of periodic paralysis, with the events in HypoPP being carbohydrate-rich meals, rest after exercise, or stress. Consequently, the first approach to disease management is life-style changes to minimize the occurrence of trigger events^3^. When these measures are insufficient, then an escalating level of pharmacological interventions is used beginning with oral K supplements, then K-sparing diuretics, and finally carbonic anhydrase inhibitors. While benefit has been established in double-blind placebo-controlled trials^4, 5^, there is an important unmet need with about 50% of patients on carbonic anhydrase inhibitors receiving inadequate control of episodic weakness or unable to tolerate these medications^6^.

The search for more effective pharmacological intervention has been driven by our understanding of the pathomechanism for episodic weakness. In periodic paralysis, the transient weakness is caused by anomalous depolarization of the muscle resting potential^1^, from a normal value of about -90 mV to the ictal range of -50 mV to -60 mV. Muscle sodium channels are inactivated by this sustained depolarization, which reduces fiber excitability and may lead to failure of action potential initiation or propagation. Interventions that favor the normal hyperpolarized resting potential of muscle are expected to be effective for reducing the risk of episodic weakness in periodic paralysis. Increasing the membrane conductance to K^+^ will hyperpolarize the resting potential. Many K^+^ channels are closed at the resting potential and drugs that promote the opening of these resting channels have been shown to hyperpolarize a wide variety of cells including neurons, skeletal muscle, smooth muscle, heart, and pancreatic beta cells^7, 8^. Two classes of so-called “K-channel openers” are in clinical use. Openers of K_ATP_ channels, such as pinacidil and diazoxide, are used to treat hypertension and hypoglycemia, respectively. Studies on biopsied muscle from patients with periodic paralysis showed pinacidil can prevent or hasten recovery from a loss of muscle force ^9, 10^, but in clinical practice intolerable hypotension occurred before a meaningful improvement of muscle function was obtained. The other clinically-used class of K channel openers promotes the activation of K_V_7 channels (also known as KCNQ channels). Retigabine (ezogabine) is a potent agonist of K_V_7.2 – K_V_7.5 channels, with the insensitive K_V_7.1 isoform expressed primarily in heart, and was available in the US as a first-in-class antiepileptic drug^11^. There have been no case reports or clinical trials of retigabine use in muscle channelopathies, but experimental studies have shown the drug reduces myotonia in mouse models of myotonia congenita^12, 13^. Neither retigabine nor any other K_V_7 agonist has previously been studied experimentally as a mechanism to ameliorate the attacks of weakness in periodic paralysis.

In this preclinical study, we tested the efficacy of retigabine to prevent the loss of force or to hasten recovery of force in a mouse model of hypokalemic periodic paralysis (HypoPP). Our knock-in mutant mouse model of HypoPP (*Scn4a* p.R669H)^14^ has a robust phenotype with paradoxical fiber depolarization and loss of contractile force in low K^+^. Susceptibility to HypoPP was determined by an ex vivo assay that measured the loss of force in response to a reduction of extracellular [K^+^] from 4.7 mM to 2 mM. Substantial protection from low-K^+^ induced loss of force was observed after pretreatment with 1 µM retigabine, and 10 µM completely suppressed any detectable loss of force. By comparison with this same assay, pinacidil was an order of magnitude less potent than retigabine for the prevention of weakness. The K_V_7-channel opener also hastened recovery. For HypoPP muscle maintained in 2 mM K^+^ the initial loss of force was reverse by application of retigabine. These data show retigabine, or other K_V_7 agonists, has the potential to reduce the severity of episodic weakness in HypoPP and is superior to previously studied K channel openers acting on K_ATP_ channels.

## MATERIALS AND METHODS

### Mouse Model of HypoPP

We previously generated a knock-in mutant mouse model of HypoPP (*Scn4a* p.R663H)^14^ homologous to the p.R669H missense mutation of the Na_V_1.4 sodium channel alpha subunit in patients with HypoPP Type 2^15^. Mutant mice have a robust HypoPP phenotype with loss of force and paradoxical depolarization of skeletal muscle in response to a challenge in low extracellular [K^+^] (< 3 mM) or to intravenous administration of insulin plus glucose^14^. Consistent with the dominant inheritance in patients, heterozygous mutant mice exhibit all the features of HypoPP. Mouse studies have revealed a gene dosage effect of the mutant allele^14^, and homozygous mutant p.R669H mice were used in this study to improve the sensitivity for detecting a beneficial effect of retigabine on the outcome measure of muscle force. The HypoPP phenotype is indistinguishable between male and female mice^14^, and the responses for soleus muscles from either sex were combined into a single group.

All procedures performed on mice were in accordance with animal protocols approved by the David Geffen School of Medicine Institutional Animal Care and Use Committee.

### Ex vivo Contraction Studies

The outcome measure for susceptibility to HypoPP was the peak isometric force of the soleus muscle, in response to a tetanic stimulation, as previously described^14, 16^. In brief, after euthanizing the animal, the soleus muscle was dissected free and suspended in a tissue bath held at 37 °C. The bicarbonate-buffered bath was bubbled continuously with 95% O_2_ / 5 % CO_2_ to maintain a pH of 7.4. The standard bath consisted of (in mM): 118 NaCl, 4.75 KCl, 1.18 MgSO_4_, 2.54 CaCl_2_, 1.18 NaH_2_PO_4_, 10 glucose, and 24.8 NaHCO_3_. The low-K^+^ solution was identical, except for the reduction of KCl to 2 mM by replacement with NaCl. The osmolality of all solutions was 290 mOsm. Retigabine (gift from Prof. M. Taglialatela, Univ. Naples) and pinacidil (Sigma-Aldrich) were prepared as 100 mM stock solutions in DMSO and diluted into the bath solution for a working concentration of 0.1 to 100 µM. The K_V_7 inhibitor, XE991 (Sigma-Aldrich), was prepared in as a 100 mM DMSO stock solution and applied to muscle in a 10 µM bath solution.

Electrical stimulation was applied by a pair of platinum wires, oriented perpendicular to the long axis of the soleus muscle. Suprathreshold stimulation (80 mA) was applied as a tetanic burst (40 pulses, 0.4 msec, at 100 Hz), under computer control. All bath solutions contained 0.25 µM D-tubocurarine (Sigma-Aldrich) to prevent any contribution to muscle excitation from motor nerve endings. Muscle force was measured with a stiff strain gain (Forte 25, World Precision Instruments) and digitally sampled at 5 KHz. Muscle contractility was monitored by measuring the peak isometric force every two minutes, and test solutions were applied by complete exchange of the bath solution.

## RESULTS

The efficacy of potassium channel openers to prevent the loss of force in HypoPP muscle was assessed by measuring isometric contractions of the soleus muscle, ex vivo, in both control (4.7 mM) and low extracellular [K^+^] (2.0 mM). The HypoPP phenotype is shown for the drug-free responses in Figure 1A. The peak isometric force during the tetanic contraction (100 Hz stimulation) was initially 11.7 gm in control [K^+^], and then decreased to 7.6 gm (35% decrease) after 4 min. in 2 mM [K^+^]. The loss of force is reversible, upon return to 4.7 mM [K^+^] (e.g., see Figure 2). For the soleus muscle from the other hindlimb of the same animal, control and low-K^+^ solutions containing 10 µM retigabine were used, and the loss of force from a 2 mM [K^+^] challenge was completely prevented (Figure 1B).

**Figure 1.**
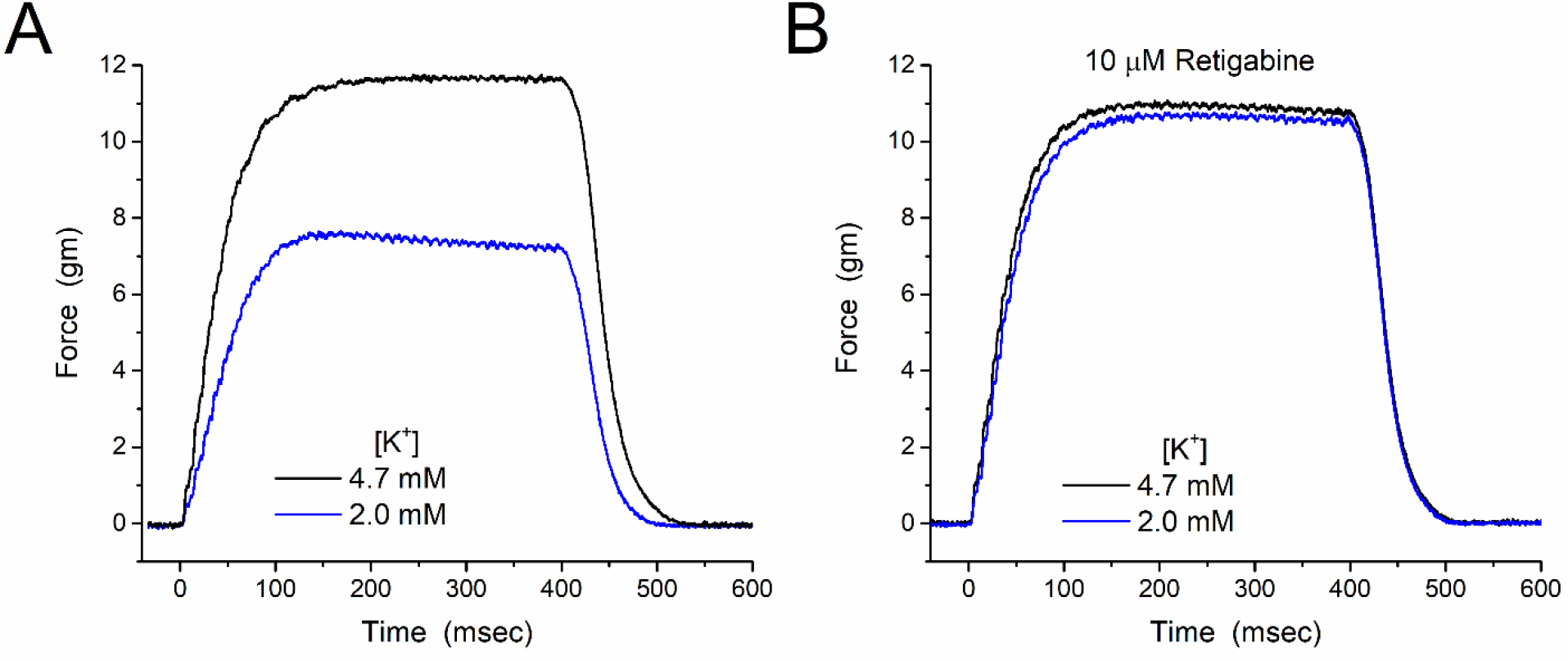
Ex vivo contraction test for susceptibility to HypoPP. (A) Isometric force measured in response to tetanic stimulation of the soleus muscle from the Na_V_1.4-R669H mouse. Baseline peak force was determined in 4.7 mM K^+^ (black line) and 4 min after bath exchange to 2 mM K^+^ (blue line). (B) The loss of force from a 2 mM K^+^ challenge was prevented in the soleus from the other hindlimb of the same animal, for which continuous exposure to 10 µM retigabine was initiated 30 min before low-K test.

**Figure 2.**
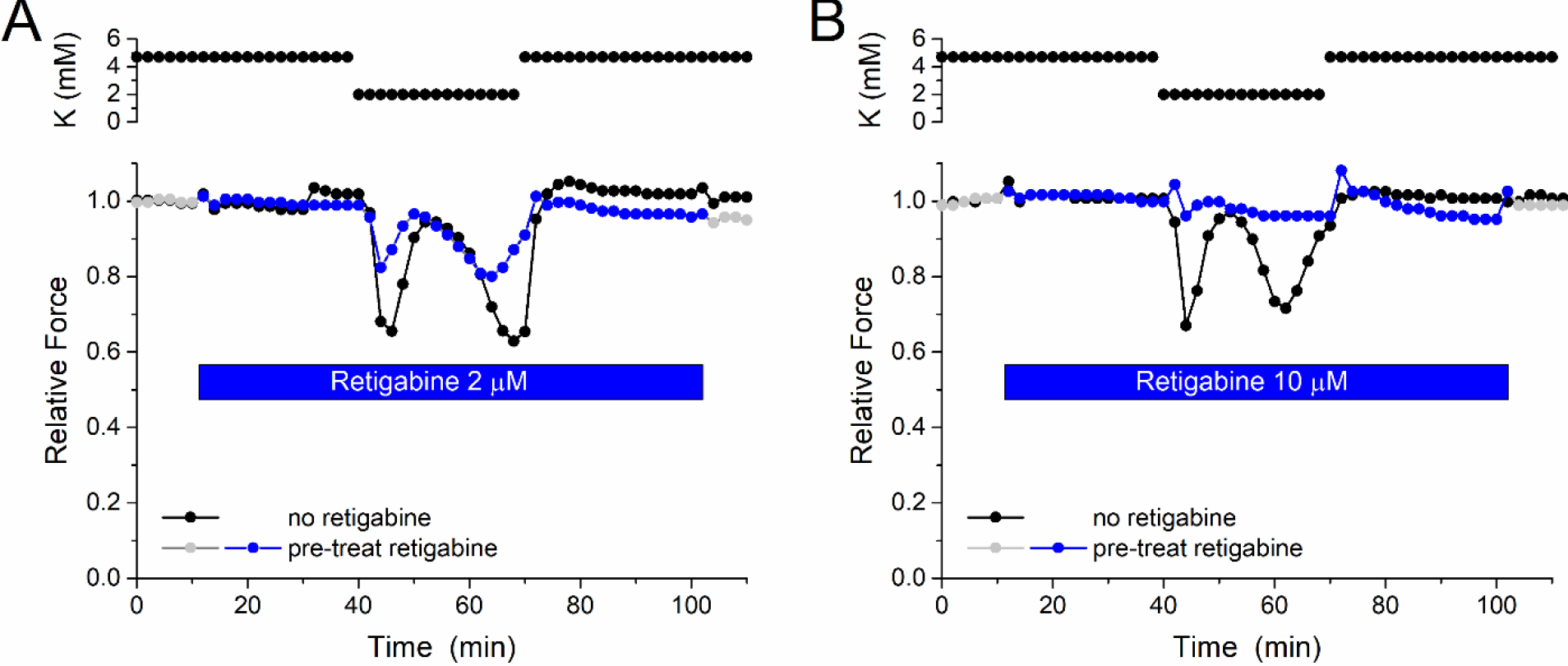
Time course of the loss of force in low K and preserved force with retigabine. (A) A tetanic contraction of the soleus muscle was recorded every 2 min, and the peak force was normalized to the average value over the first 5 measurements in 4.7 mM K^+^. During a 30 min challenge with 2 mM K^+^ (40 min to 70 min) the peak force waxed and waned in an oscillatory pattern with a nadir of 40% loss (black trace). Pre-treatment with 2 µM retigabine markedly attenuated the loss of peak force. (B) A similar protocol with 10 µM retigabine completely prevented the loss of force in a 2 mM K^+^ challenge. The paired responses in (A) are from the soleus muscles of the same Na_V_1.4-R669H mouse, while those in (B) are paired responses from another Na_V_1.4-R669H mouse.

The time course of the relative force in response to a 30 min. low-K^+^ challenge, with and without pretreatment with retigabine, is shown in Figure 2. The relative force was stable before the low-K^+^ challenge and was not affected by the addition of retigabine (time = 10 min. in Figure 2) over the range of concentrations used (0.1 to 20 µM). As we previously reported^14^, the loss of force during a low-K^+^ challenge in the Na_V_1.4-R669H HypoPP mouse in drug-free conditions may exhibit oscillations with spontaneous periods of recovery for the soleus muscle. While this oscillatory response is variable across different mice, the concordance is high between the left and right soleus muscles of a single mouse, as shown by the in-phase oscillations for control and retigabine in Figure 2A or for paired drug-free recordings in Wu et al. ^14^. The representative responses in Figure 2B reaffirm that pre-treatment with 10 µM retigabine completely prevents the loss of force in 2 mM [K^+^], whereas partial protection occurs with 2 µM of the drug.

Our method to quantify the extent of protection from loss of force by retigabine is illustrated in Figure 3. This example was selected to illustrate the maximum variability in the extent of protection within a single trial (compare the responses at 24 min to those at 40 min.). Qualitatively, even 1 µM retigabine was sufficient to provide substantial protection. The relative protection was calculated from the responses for each pair of soleus muscles from the same mouse serving as an internal control. In almost every trial (22 out of 24), two clear minima occurred during the 30 min. low-K^+^ challenge. The relative protection for the retigabine response, defined as 1- (force loss in retigabine)/(force loss in control) was calculated for each of the two minima, and the average value was used as the quantitative estimate for the retigabine protection in that trial. Each soleus muscle was used for only a single trial (i.e. control or retigabine) because cumulative effects and run-down may occur if this long protocol (typically 70 min.) was repeated.

**Figure 3.**
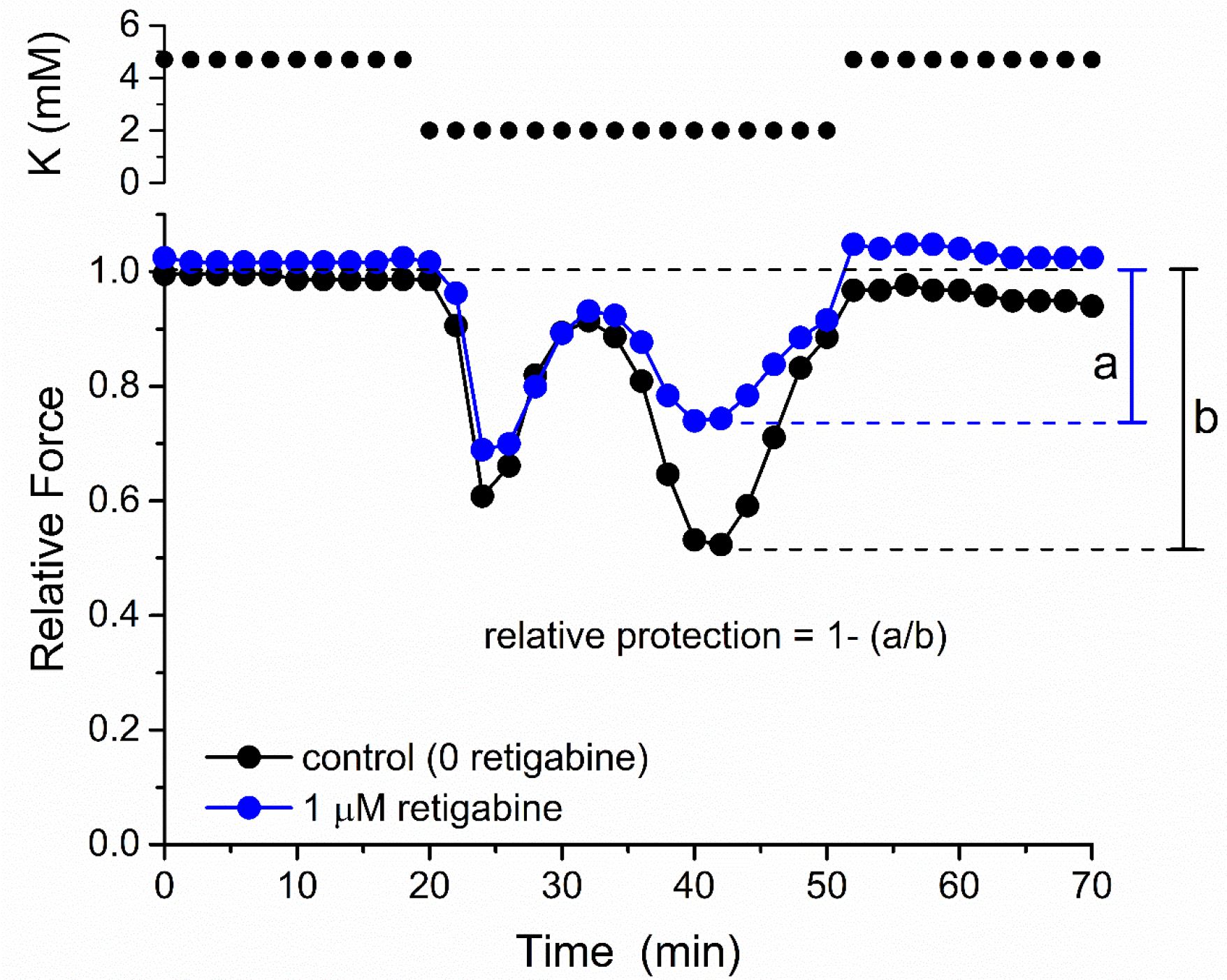
Protocol to quantify protection from loss of force. This representative contraction test shows variability in the extent of protection by retigabine; compare the small difference in control (black) to retigabine (blue) nadirs at 24 min. versus the larger difference at 42 min. In general, the protection in 1 µM retigabine was less than for 2 µM or 10 µM (Figure 2). The extent of protection was measured at each nadir (generally two within the 30 min low-K challenge) as the ratio of the loss of force in retigabine (distance “a”) to the loss in control with no drug (distance “b”). The average of these two measurements was used as the quantitative estimate for the extent of protection for that animal. The low-K^+^ challenge (2 mM) was applied from 20 to 50 min.

The dose-response relation for the relative protection from loss of force by pre-treatment with retigabine is shown in Figure 4. The concentration-dependent protection had an equivalent *K*_*d*_ = 0.82 ± 0.13 µM, based on the best-fit by a single binding-site model. A limited series of measurements were also performed with pinacidil, a K_ATP_ channel opener, that has previously been shown to protect against loss of force for an ex vivo contraction assay of human HypoPP muscle at a concentration of 100 µM^9^. Using our Na_V_1.4-R669H mouse model, we confirmed that pre-treatment with pinacidil protects HypoPP muscle from a loss of force in a low-K^+^ challenge (Figure 4), however the potency was about 10-fold lower (*K*_*d*_ = 8.1 ± 0.5 µM) compared to retigabine.

**Figure 4.**
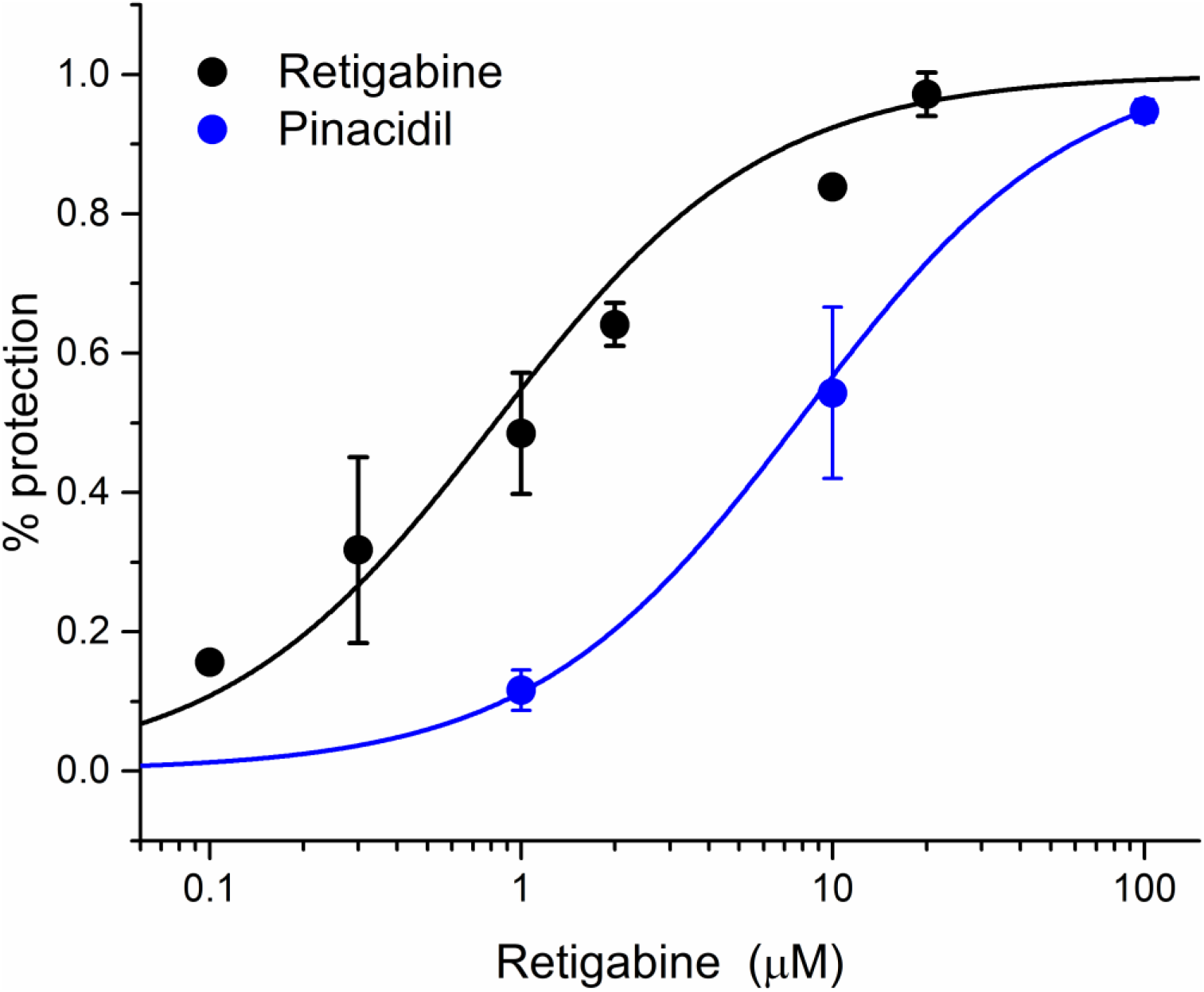
Dose-Response relation for the protection by retigabine and by pinacidil. Each symbol is a mean value for the relative protection of force in pair-wise comparisons of drug to no drug for a challenge with 2 mM K^+^ (same mouse, two different soleus muscles). The curves show fitted estimates for a single binding-site model with *K*_*d*_values of 0.82 ± 0.13 µM for retigabine and 8.1 ± 0.49 µM for pinacidil. The therapeutic range for unbound drug concentration is shown by the shaded region.

We also tested whether application of retigabine would hasten the recovery from a low-K induced loss of force in HypoPP muscle. Two examples from soleus muscles of separate HypoPP mice are shown in Fig 5. A 2 mM K^+^ challenge was applied first, and then after confirming a loss of peak force, the bath was exchanged by a new 2 K^+^ solution that also contained 10 µM retigabine. Application of retigabine at 10 min. after the onset of reduced force (Fig 5A) or even after 30 min when two loss-of-force cycles were complete (Fig. 5B), produced a sustained improvement in force and prevented any further cyclical reduction of force. The improved level of tetanic force at about 83% of baseline in 4.7 K^+^ is comparable to the performance of WT soleus in 2 mM K^+^ of 89% (Fig. 5, red lines).

**Figure 5.**
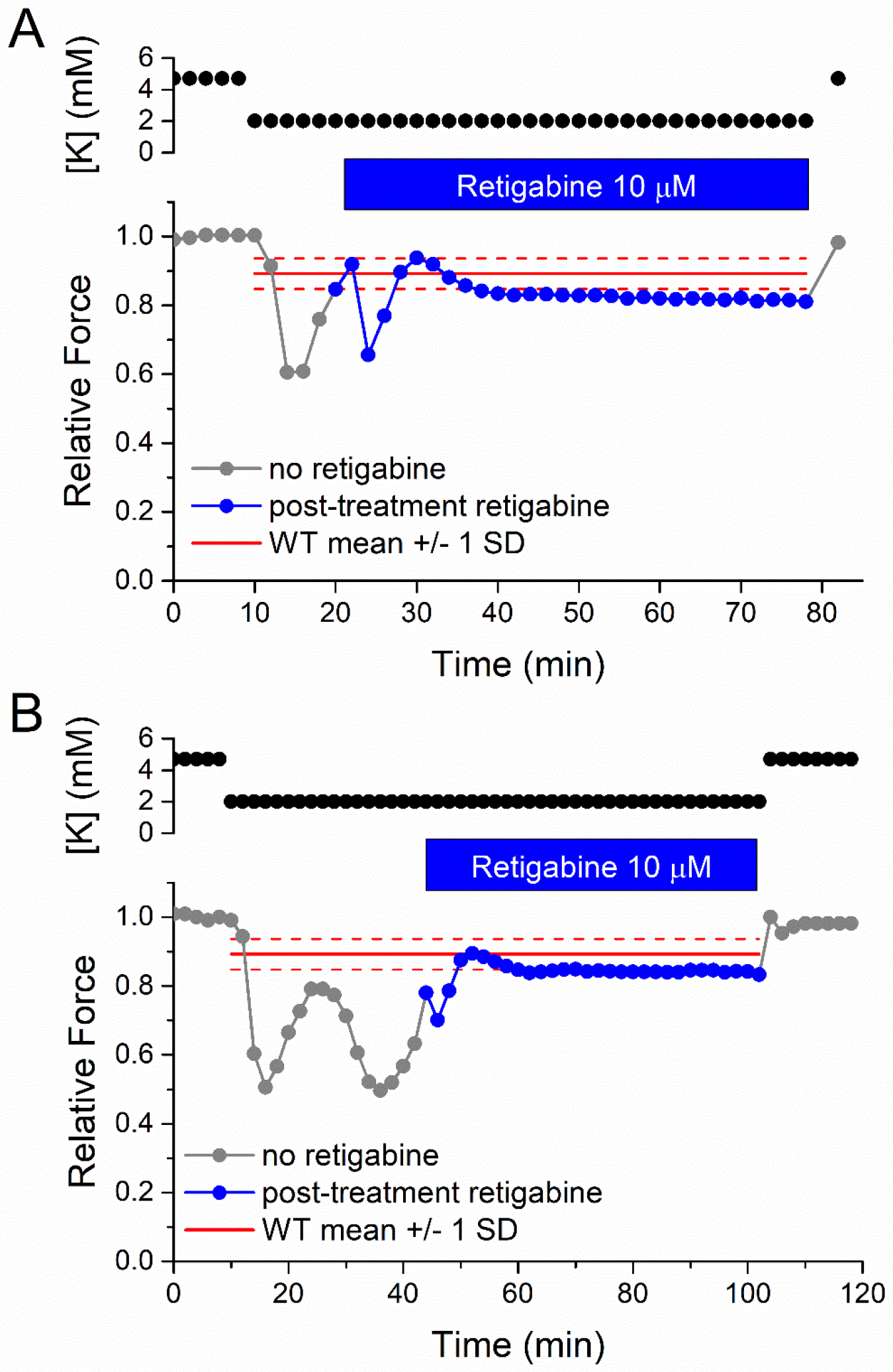
Retigabine promotes recovery from the loss of force for HypoPP muscle while in low K^+^. Examples of retigabine-induced rescue from loss of force are shown for two representative experiments. (A) 10 min after the onset of 2 mM K^+^, the oscillatory loss of force is aborted by application of retigabine. (b) 30 min after 2 mM K^+^ with two cycles of reduced force, retigabine restores the relative force to a sustained value of 0.83. In both panels, the red line shows the force mean ± 1 SD for WT soleus in 2 mM K^+^.^14^

To confirm the beneficial effects of retigabine were dependent upon currents conducted by K_V_7 – type K^+^ channels, we assessed the effect of a K_V_7 inhibitor, XE991^17^. As shown in Fig. 6, exposure to 10 µM XE991 did not alter the peak tetanic force in 4.7 mM K^+^ (10 to 20 min, magenta symbols). The other soleus muscle from the same HypoPP mouse was not exposed to XE991. Retigabine was then applied to both soleus muscle preparations, followed by a low-K challenge. A marked loss of force in low-K occurred for the XE991 + retigabine exposed soleus, whereas the paired HypoPP muscle was protected by retigabine. These data show the beneficial effect of retigabine is blocked by XE991 and suggest that basal activity of K_V_7 channels limits the severity of the loss of force in low K^+^.

**Figure 6.**
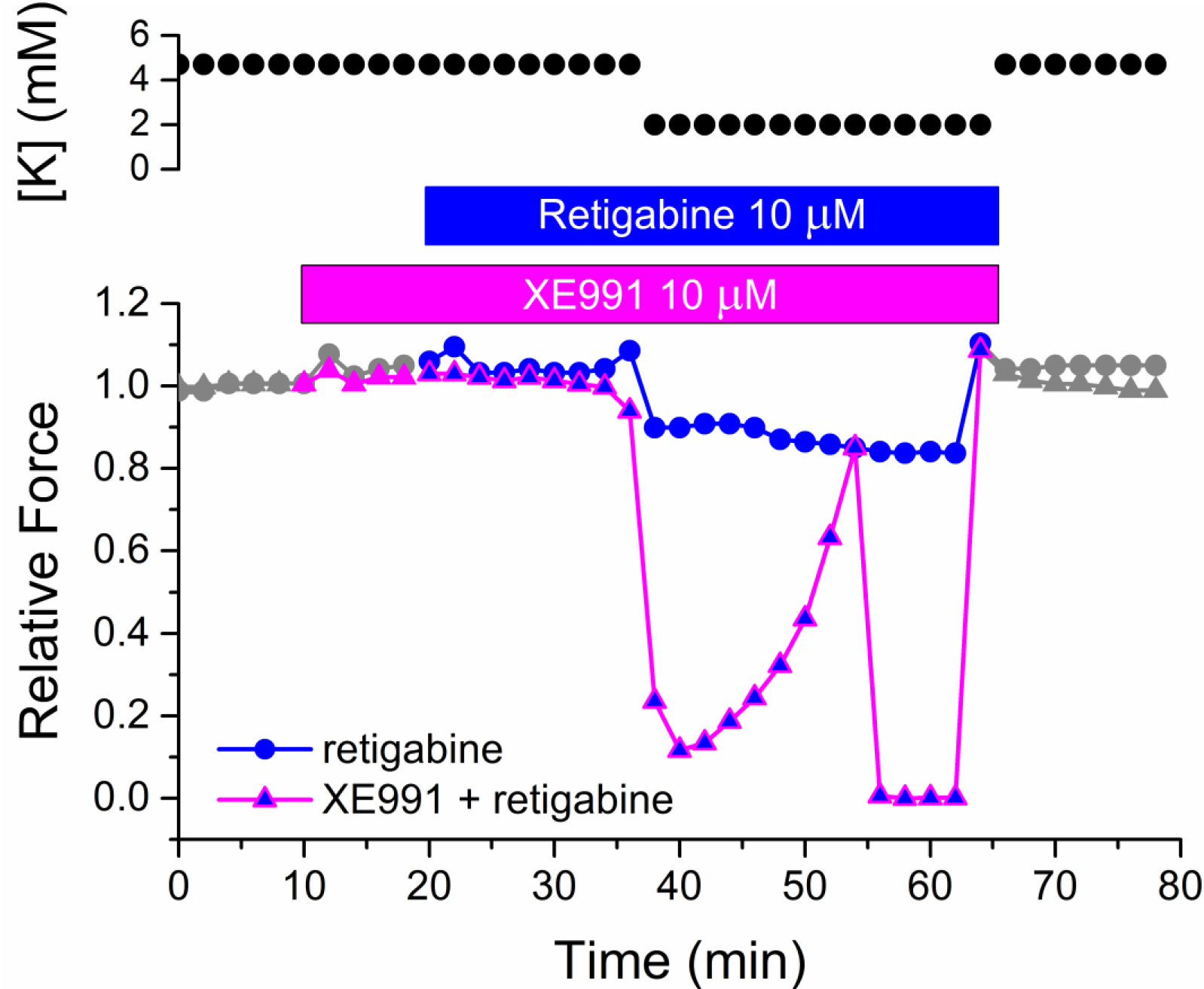
The protective effect of retigabine was prevented by the K_V_7 channel inhibitor XE991. Paired soleus muscles from the same HypoPP mouse were used to test for an effect of XE991. Application of XE991 to one muscle did not alter the baseline peak tetanic force (10 min., magenta symbols). Pretreatment with retigabine at 20 min (blue symbols) prevented the loss of force during a low-K challenge (40 to 60 min.). In contrast, an exceptionally large loss of force occurred for the XE991-exposed soleus muscle, despite the same pretreatment with retigabine.

## DISCUSSION

The therapeutic potential of K^+^ channel openers to reduce the frequency and severity for the attacks of weakness in periodic paralysis has been recognized for over 30 years^10^. The fundamental principle is that because increasing the membrane conductance to K^+^ will hyperpolarize the membrane potential toward the Nernst potential for K^+^ (*E*_*K*_ ≈ -95 mV), this class of drugs will counteract the sustained anomalous depolarization of the resting potential that causes transient weakness in periodic paralysis by inactivation of sodium channels. Many criteria must be met, however, for this strategy to be safe and effective. For example, many K^+^ channels are activated by depolarization, and the drug-induced augmentation of channel opening must occur over an operational voltage range that is relevant for stabilization of a normal resting potential (-80 to -95 mV) and extend to the anomalous depolarization during paralysis (typically - 50 to -60 mV). This voltage-dependent property will determine whether the increased K^+^ conductance will primarily affect the resting potential, the action potential waveform (e.g. amplitude and duration), or both. Another concern is the expression pattern across different tissues for the K^+^ channels that are activated by the drug; expression in skeletal muscle for the desired effect, with lower expression in other tissues to minimize side effects. The drug potency (small dissociation constant, *K*_*d*_) and magnitude of the K^+^ conductance increase are critical as well. If the conductance increase is very small, then the drug may not produce a meaningful reduction in the susceptibility to attacks of weakness. Conversely, a very large K^+^ conductance increase may impair contractility by reducing muscle fiber excitability.

The effectiveness of retigabine at low µM concentrations to protect HypoPP muscle from a loss of force in a low-K^+^ challenge (2 mM) demonstrates the therapeutic potential of K_V_7.x channel openers in the symptomatic management of HypoPP. The K_V_7.1 – K_V_7.5 family of voltage-gated K^+^ channels is encoded by *KCNQ1*-*KCNQ5* genes, so-named because the founding member, *KCNQ1*, was identified as a disease locus for type 1 long QT syndrome (LQT1)^18^. The predominant K_V_7 channel in the heart is K_V_7.1, and importantly, this isoform is 100-fold less sensitive to retigabine than K_V_7.2-K_V_7.5^19^, which explains the absence of cardiac side effects. The K_V_7.2 - K_V_7.5 isoforms were initially characterized as “neuronal”, with K_V_7.2, K_V_7.3, K_V_7.5 expressed in the central nervous system, K_V_7.4 in the cochlea, and K_V_7.2 in the peripheral nervous system. Skeletal muscle was not initially considered to be a site of significant K_V_7.x expression, but subsequent studies using mouse and human skeletal muscle have detected K_V_7.1-K_V_7.5 transcripts by RT-PCR, and K_V_7.2-K_V_7.4 subunit protein by both immunoblot and immunohistochemistry^20^. Changes in the expression pattern of K_V_7.x subunits with muscle proliferation and differentiation have implicated a role in development.

A physiological role has not yet been established for the voltage-gated K^+^ current conducted by K_V_7 channels in skeletal muscle. Voltage-clamp studies to characterize K_V_7.x currents in muscle have not been reported, most likely because of the technical difficulty in isolating this component of the K^+^ current from the contributions by a multitude of K^+^ channels in this tissue (over 20 described). The beneficial effect of retigabine is completely prevented by XE991 (Fig. 6), which at 10 µM is a specific inhibitor of K_V_7 channels^21^ and supports our interpretation that the K channel-opening action of retigabine is the mechanism of drug action. Because low-dose retigabine protects HypoPP muscle from a low-K^+^ induced loss of force and also suppresses after-discharges in mouse models of myotonia (using pharmacologic block of the chloride conductance^13^ or by disruption of the *Clcn1* gene^12^), the K_V_7.x K^+^ current clearly has an important role in regulating muscle excitability. Prior experiments (using current-clamp) to explore the mechanism for retigabine-induced suppression of myotonia did not detect a change in fiber input resistance, resting potential or action potential properties of mouse muscle upon exposure to 20 µM retigabine^12^. While disappointing, this failure is not surprising because it is well known that almost imperceptible perturbations in channel behavior can have profound effects on excitability in muscle channelopathies^1^. Electrophysiological approaches with greater sensitivity and specificity will be required to elucidate the normal role of K_V_7.x K^+^ currents in muscle and the beneficial effect of activation by retigabine.

While the chronic use of retigabine as a prophylactic antiepileptic drug for refractory partial-onset seizures was discontinued in the US in 2017 because of adverse effects (primarily skin discoloration^22^ and also CNS effects^23^ with dose-dependent dizziness, somnolence, headache), the enthusiasm for K_V_7 channel activators remains strong for management of epilepsy (especially KCNQ2 developmental and epileptic encephalopathies), pain management, for neuroprotection in degenerative diseases, and for mood stabilization. Derivatives of retigabine with reduced dimer formation and therefore lack skin discoloration are in clinical trials for KCNQ2-DDE (XEN496 in children; NCT04912856) and for focal epilepsy in adults (XEN1101; NCT03796962). Another KV7 opener, GRT-X, that is chemically unrelated to retigabine is in development. These pipeline drugs also act by activation of K_V_7 channels, and therefore are likely to be effective in ameliorating attacks of periodic paralysis.

Moreover, our data showing rescue of contractility (Fig. 5) demonstrates the potential for short-term administration of retigabine, or newer K_V_7 openers, as abortive therapy for an acute attack of weakness in HypoPP. After a single 100 mg oral dose, a *C*_*max*_ of 390 ng/ml occurs within 1.5 hrs., and the half-life is 8 hrs^24^. An unbound retigabine fraction of 0.9 µM will be achieved with a single 400 mg dose (assumes 80% protein binding). Based on this observation, a clinical trial for efficacy of K_V_7 openers could be performed by assessing the rate of recovery of the compound muscle action potential in the long exercise test for periodic paralysis, as was done to studies of bumetanide^3^. The functional defect in periodic paralysis operates like a binary switch, with a sustained shift between the normal *V*_*rest*_ and an anomalously depolarized value. Consequently, a single dose of retigabine may be sufficient to “reset” *V*_*rest*_ to the normal range and suppress a recurrent attack of weakness over the minutes to hours required for correction of extracellular [K^+^]. An adverse effect of retigabine on skeletal muscle function is unlikely. One report described a dose-dependent reduction of contractile force during a 3 sec tetanic stimulation of rat muscle, with an IC_50_ of 1 µM, presumably from a reduced fiber excitability^25^. Our studies with up to 20 µM retigabine show no impairment of peak isometric force during a 400 msec tetanic contraction, applied once every two minutes for two hrs. Similarly, the prior two studies on the effect of retigabine in mouse models of myotonia did not report a reduction of ex vivo tetanic force with 20 or 30 µM retigabine^12, 13^. In vivo mouse and rat studies that used a very high dose of retigabine (30 mg/kg which is 5.7 times larger than the maximal oral dose of 400 mg for a 70 kg human) showed a loss of force with tetanic contractions lasting seconds^12, 25^, but this result is not likely to be relevant to patient care. In our opinion, the “fatigue” described as an adverse effect of retigabine in AED trials is more likely to be of central origin rather than muscle fiber dysfunction.

## Acknowledgements

This work was supported by the National Institutes of Arthritis, Musculoskeletal, and Skin diseases (R01AR063182 and R01AR078198).

## Author Contributions

MQ performed the experiments, analyzed the data, and edited the manuscript. SC designed the experiments, analyzed the data, and quote the manuscript.

